# Optimization-based dietary recommendations for healthy eating

**DOI:** 10.1101/2024.12.17.628891

**Authors:** Xu-Wen Wang, Scott T. Weiss, Frank B. Hu, Yang-Yu Liu

## Abstract

Various diet scores have been developed to assess compliance with dietary guidelines. Yet, enhancing those diet scores is very challenging. Here, we tackle this issue by formalizing an optimization problem and solving it with simulated annealing. Our optimization-based dietary recommendation (ODR) approach, evaluated using Diet-Microbiome Association study data, provides efficient and reasonable recommendations for different diet scores. ODR has the potential to enhance nutritional counseling and promote dietary adherence for healthy eating.

## INTRODUCTION

Diet plays a critical role in the development of many chronic diseases^1,2^. To mitigate the risk of these ailments, many countries have developed national dietary guidelines to provide context-specific advice and principles on healthy diets and lifestyles. Various diet scores have been proposed to quantify adherence to those dietary guidelines or specific dietary patterns. For example, the Healthy Eating Index (HEI)^3^ was developed by the United States Department of Agriculture to measure adherence to the Dietary Guidelines for Americans. HEI evaluates the diet quality based on the intake of food groups such as fruits, vegetables, whole grains, dairy, and protein, and limits components like saturated fat, sodium, and added sugars. HEI has been updated over the years to reflect changes in dietary guidelines. The Alternative Healthy Eating Index (AHEI)^4^ is similar to the HEI but tailored to better reflect associations with chronic disease prevention, such as lowering the risk of cardiovascular disease, diabetes, and cancer. It emphasizes plant-based foods, healthy fats, and limits red meat, sugary drinks, and trans fats. Mediterranean Diet Score (MDS)^5^ was developed to quantify adherence to the Mediterranean Diet. MDS evaluates the consumption of typical Mediterranean foods based on the intake of nine items: vegetables, legumes, fruit and nuts, dairy, cereals, meat and meat products, fish, alcohol, and the ratio of monounsaturated to saturated fat. The Alternate Mediterranean diet score (AMED)^6^ was adapted to better align with the dietary habits and food availability in non-Mediterranean regions, particularly the U.S. The Dietary Inflammatory Index (DII) was designed to evaluate the inflammatory potential of an individual’s diet by considering the effect of each of the 45 ‘food parameters’ on six inflammatory biomarkers: IL-1b, IL-4, IL-6, IL-10, TNF-a and C-reactive protein^7^. The forty-five pro- and anti-inflammatory ‘food parameters’ consist of whole foods, nutrients, and other bioactive compounds derived from the literature review. In order to standardize an individual’s dietary intakes, a global composite database, which comprises eleven food consumption data sets from various countries, was utilized to express individuals’ dietary intakes relative to the range of intakes of the forty-five food parameters observed among these diverse populations.

Those diet scores are useful tools in both clinical and research settings to assess the quality of an individual’s diet and to evaluate the impact of dietary patterns on health outcomes. Yet, mathematically optimizing those diet scores could be a challenging task. This is because those scores are typically computed from the sum of multiple food and nutrition items, and increasing the intake of one food or nutrition item might affect the intake of other items defined in the diet score in two different ways. First, consuming more of one food group can reduce the consumption of other groups due to limits in total caloric intake or food volume capacity. This is known as *dietary displacement*^8^. Second, there are *interdependencies* between food and nutrient components in many diet scores. Consider HEI2015 as an example^9^. HEI2015 is a diet score based on criteria in the Technical Report of the Healthy Eating Index-2015. HEI2015 includes 13 energy-adjusted components representing the major food groups in recommendations from the 2015 Dietary Guidelines for Americans: 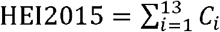, where *C*_*i*_ is the score of each component-i in HEI2015. Intakes between the minimum and maximum standards are scored proportionately in the calculation of *C*_*i*_. We notice that four components (i.e., saturated fat, sodium, fatty acids, and sugars) in the HEI2015 computation are derived from the food amount. Thus, increasing the serving of those food components might reduce the scores of these nutrition components, and this interdependency or trade-off renders the optimization of HEI2015 a very challenging task. For instance, an increase (or decrease) in the “total vegetable” component is sometimes associated with a decrease (or increase) in HEI2015 (**Fig.1a**). A similar pattern is observed for DII, where changes in DII may not align consistently with changes in specific components in DII, such as vitamins B12 and B6 (**Fig.1b**).

**Figure 1:**
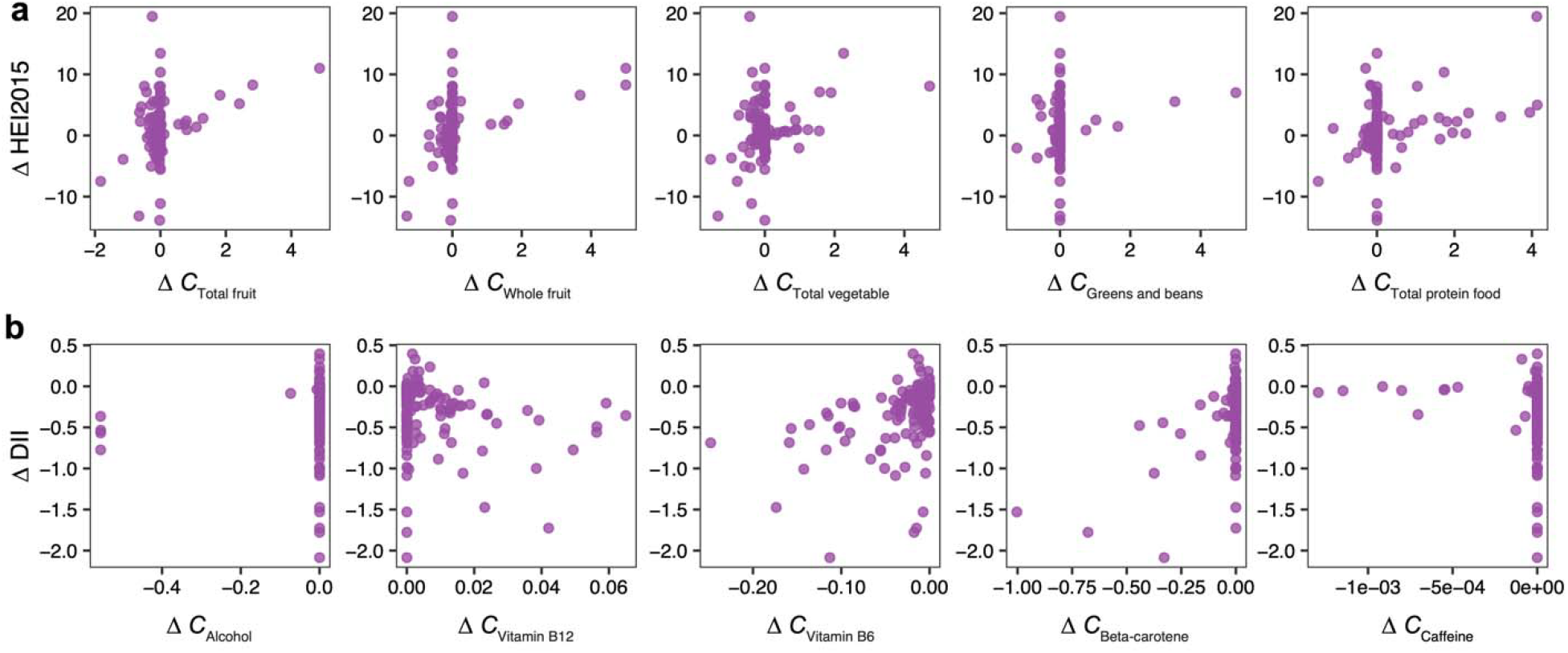
Optimizing diet scores is nontrivial because of the interdependency of food and nutrition components in the diet scores. Correlation between the change in the overall diet score and the change of the first five components (columns) in HEI2015 (a) and DII (b). Each panel shows the change of the overall diet score and the change of a component score after randomly perturbing the dietary intake of a subject in DMAS by adding a new food item. We performed 200 perturbations. For each perturbation, we randomly selected a subject on a random day from DMAS and calculated his/her original diet score along with the score for each component in the diet score. Next, we randomly added a new food item (selected from the pooled list of all food items consumed by all subjects from DMAS) with the gram weight of the added food item matching the food consumption pattern in DMAS.

Here, we formalized the diet recommendation as an optimization problem by considering the diet score as the target function and optimizing it using the simulated annealing (SA) algorithm^10,11^. We evaluated this optimization-based dietary recommendation (ODR) approach using real data^12^, demonstrating its potential for personalized dietary recommendations.

## RESULTS

### Formalization of the diet recommendation problem

As we mentioned above, there are many diet scores to quantify adherence to dietary guidelines or patterns. Our proposed algorithm provides a universal framework for personalized diet recommendations to optimize any diet scores. For a given food profile ***f***= (*f*_*l*_, *f*_*2*_,*…, f*_*N*_) of a subject collected from a dietary assessment tool, e.g., Automated Self-Administered 24-hour (ASA24), we can compute its nutrient profile ***q*** = (*q* _*l*_, *q*_*2*_, *…, q*_*M*_) based on a food composition database, e.g., Frida^13^, USDA’s Food and Nutrient Database for Dietary Studies^14^, and the Harvard food composition database^15^. A diet score S can be expressed as a function of its food profile 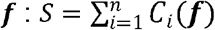. Here, *C*_*i*_ (***f***) represents the i-th component in the diet score, which is a typically binary function of the food intake profile. n is the total number of components in the diet score. Our goal is to maximize S by recommending an optimal food profile.

### Optimization-based dietary recommendation (ODR)

To maximize the diet score *S*, we used simulated annealing, which is a classical optimization method inspired by the annealing process in metallurgy. Simulated annealing aims to find a global minimum of a function by allowing occasional acceptance of worse solutions to escape local minima. The algorithm starts with a high “temperature” that permits greater exploration of the solution space by accepting both better and worse solutions. As the temperature gradually decreases, the algorithm becomes more selective, favoring improvements and reducing the likelihood of accepting inferior solutions. This balance between exploration and exploitation enables simulated annealing to navigate complex, multimodal optimization landscapes effectively. The detailed procedure of maximizing the diet score via simulated annealing is shown in **Methods**. To ensure that the recommendations align with a practical meal plan, we focused the food items pool from a real dataset --- the Diet-microbiome association study (DMAS)^12^, and limited the number of food items for each of the eight eating occasions (i.e., breakfast, brunch, lunch, dinner, supper, just a drink, snack, and other) to a reasonable range. To ensure an individual’s dietary pattern remains consistent, we required that at least half of the recommended food items match those in the original diet. Finally, each recommended food item was assigned to one of the eight eating occasions.

### Demonstration of ODR using real data

To demonstrate the power of the ODR approach, we apply it to the DMAS dataset^12^, consisting of 24-h food records from 34 healthy human subjects collected daily over 17 days. We first examined the total food amount of each subject across different days, finding that despite the total food amount of each subject showing variation, the mean food amount among different subjects across different days remains conserved, i.e., displaying relatively small variation around 3,000 grams (see **Fig.S1a**). This variation among food profiles is reflected in HEI2015, which also shows variation across days (see **Fig.S1b**). However, we found that the mean HEI2015 among different subjects is stabilized at 60, implying that the diet quality of most subjects is moderate. Then, we applied ODR to the dietary record with the lowest HEI2015 originally among 30 subjects over 17 days (**Fig.2a**) by choosing distance parameters *r* = 0.4. We found that the HEI2015 increased from 26 to 76, and some unhealthy items, for instance, refined grains, chips, and popcorn, were reduced, but some healthy items, for instance, dairy and fruits were increased. Certain healthy items in the original diet, such as milk and yogurt, were kept in the recommended diet (**Fig.2a**). This alignment between changes in the HEI2015 components and food gram adjustments suggests that the SA recommendations are reasonable.

**Figure 2:**
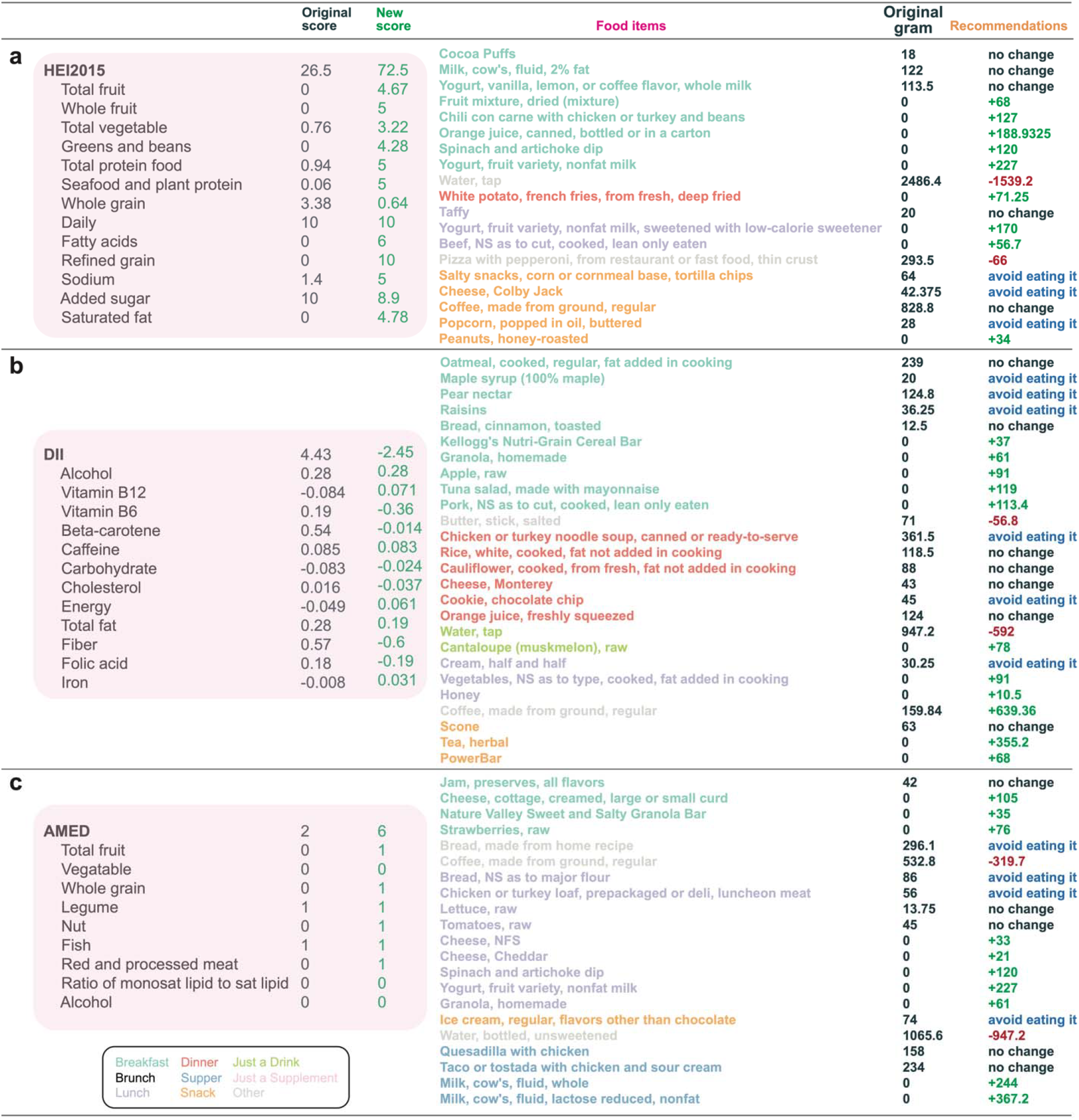
Optimizing three different diet scores yield different dietary recommendations. Diet scores: HEI2015 (**a**), DII (**b**), and AMED (**c**). For each diet score, we show the original and new overall score, as well as the original and new component scores. We also show the grams of the original food items in the dietary intake (black) and the recommendations made by the optimization algorithm (with the parameter food profile distance). Each food item was colored by its corresponding mealtime. For DII, for visualization purposes, we only listed the top 12 components among its 45 components (or ‘food parameters’).

In addition to optimizing the HEI2015, we also demonstrated that SA could make reasonable recommendations for the anti-inflammatory diet by decreasing DII (**Fig.2b**). We found that the DII decreased from 4.7 to −2.5, and some pro-inflammatory items, for instance, butter (with saturated fat), cookies (refined grains and added sugars), and rice (refined grains) were reduced, but some anti-inflammatory items, e.g., vegetables (fiber), apple (fiber), tune (antioxidants and vitamins) and tea (polyphenols and flavonoids) were increased. Certain anti-inflammatory items in the original diet, such as oatmeal (fiber) and cauliflower (antioxidants and fiber), were kept in the recommended diet (**Fig.2b**).

Finally, we demonstrated SA in optimizing the AMED score. Although all components in AMED are food-related, making the recommendations appear straightforward, ODR is still capable of generating recommendations in an automated manner. We found that the AMED increased from 2 to 6, with the components, such as whole grains, nuts, vegetables, and meat, each contributing one additional point (**Fig.2c**). We then compared the gram amounts of each food in the original diet with those in the recommended diet, observing that certain unhealthy refined and processed foods, such as bread, chicken loaf, and ice crease were reduced. Conversely, certain healthy foods, such as dairy, fresh produce, and grain, were added to the recommended diet, contributing to the nuts and whole grains categories, respectively. In addition, certain healthy foods in the original diet, e.g., raw tomato and lettuce, were kept in the recommended profile (**Fig.2c**).

## DISCUSSION

In this work, we formalized the prescription of dietary recommendations as an optimization problem and solved it using the classical simulated annealing algorithm. This optimization-based dietary recommendation approach can provide tailored recommendations designed to optimize a given diet score, offering specific guidance on food choices to enhance alignment with the chosen dietary pattern (e.g., Mediterranean diet) or dietary guidelines, thereby promoting continuous improvement in dietary quality.

This innovative application of optimization algorithms will allow us to create a highly adaptable model that personalizes dietary advice based on the complex interactions between various dietary components and individual characteristics. By maximizing the diet score through this optimization process, we ensure that individuals receive dietary guidance that is not only evidence-based but also mathematically optimized to enhance adherence to a desired dietary guideline or pattern. To our knowledge, this study represents the first application of a statistical physics-based optimization in the field of personalized nutrition and dietary adherence. This allows us to model dietary recommendations in a novel, data-driven manner, which opens new frontiers for personalized health interventions. By leveraging statistical physics, we can navigate the complex, multidimensional nature of diet adherence, integrating nutritional science with advanced computational methods to provide precise, personalized feedback. This methodological innovation holds the potential to revolutionize dietary adherence strategies in nutritional research, providing a scalable and scientifically rigorous tool that can be adapted for various dietary patterns and health outcomes.

## Methods

### Data source

The Diet-Microbiome Association Study (DMAS) is a longitudinal investigation involving 34 healthy participants. Over 17 consecutive days, daily dietary intake data and stool samples were collected. Dietary intake was assessed using the Automated Self-Administered 24-hour (ASA24) dietary assessment tool. The nutritional composition of foods in DMAS was analyzed using ASA24-2016, which assigns nutrient values to foods based on the USDA Food and Nutrient Database for Dietary Studies (FNDDS 2011–2012). All healthy eating scores were computed using the R package “dietary index”^16^ without summarizing the food group and nutrient intake over all days reported per individual per day.

### Optimizing diet scores using simulated annealing

Given an initial food intake profile 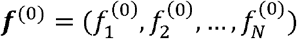 and its corresponding diet score *S*_0_, the optimal dietary recommendation to maximize the diet score is obtained by the following steps: (1) Randomly perturb the current food profile ***f***^(c)^ by three possible strategies with an equal probability to yield a new food profile ***f***^(n)^: (i) randomly replace one of the present food item with another absent food item. (ii) add a new food item, and the gram of this food is drawn from the DMAS^12^. (iii) remove one of the present items. (2) Compute the new diet score *S*^(n)^. (3) Update the current food profile ***f***^(c)^ to ***f***^(n)^ if *S*^(n)^ > *S*^(c)^ or with probability *p* = exp [*S*^(n)^ − *S*^(c)^)/*T*] if *S*^(n)^ < *S*^(c)^. Repeat steps (1)-(3) 200 times to obtain the final recommendation *f*^(r)^. During each step *t*, the “temperature” *T* is decreased geometrically^17^ as 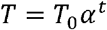, where the initial temperature *T*_0_ is set to be 500, and = 0.999. We will also introduce a hyperparameter r to control the total gram dissimilarity between the original food profile and the recommended food profile *d*(*f*^(0)^, *f*^(r)^) < *r*, where *d*(f^(0)^, *f*^(r)^)= ∣ ∑*f*^(0)^ − ∑*f*^(r)^I∣ / ∑*f*^(0)^ is the relative distance between the two food profiles. Larger parameter r allows for larger perturbations to the current dietary intake and yields a higher diet score (see **Fig.S2**).

## Acknowledgements

We acknowledge grants from the National Institutes of Health (R01AI141529, R01HD093761, R35CA253185, RF1AG067744, UH3OD023268, U19AI095219, U01HL089856, U01-152905, and U01-167552) and Cancer Grand Challenges Team PROSPECT. Y.-Y.L. acknowledges funding support from the Office of the Assistant Secretary of Defense for Health Affairs, through the Traumatic Brain Injury and Psychological Health Research Program (Focused Program Award) under award no. W81XWH-22-S-TBIPH2, endorsed by the Department of Defense. X.-W.W. acknowledges the funding support from National Institutes of Health (K25HL166208).

## Author Contributions

Y.Y.L. conceived and designed the project. X.W.W. performed all the numerical calculations. X.W.W. and Y.Y.L. wrote the manuscript. All authors analyzed the results, edited, and approved the manuscript.

## Data accessibility

The data and code used in this work are available at https://github.com/spxuw/ODR.

**Figure S1:**
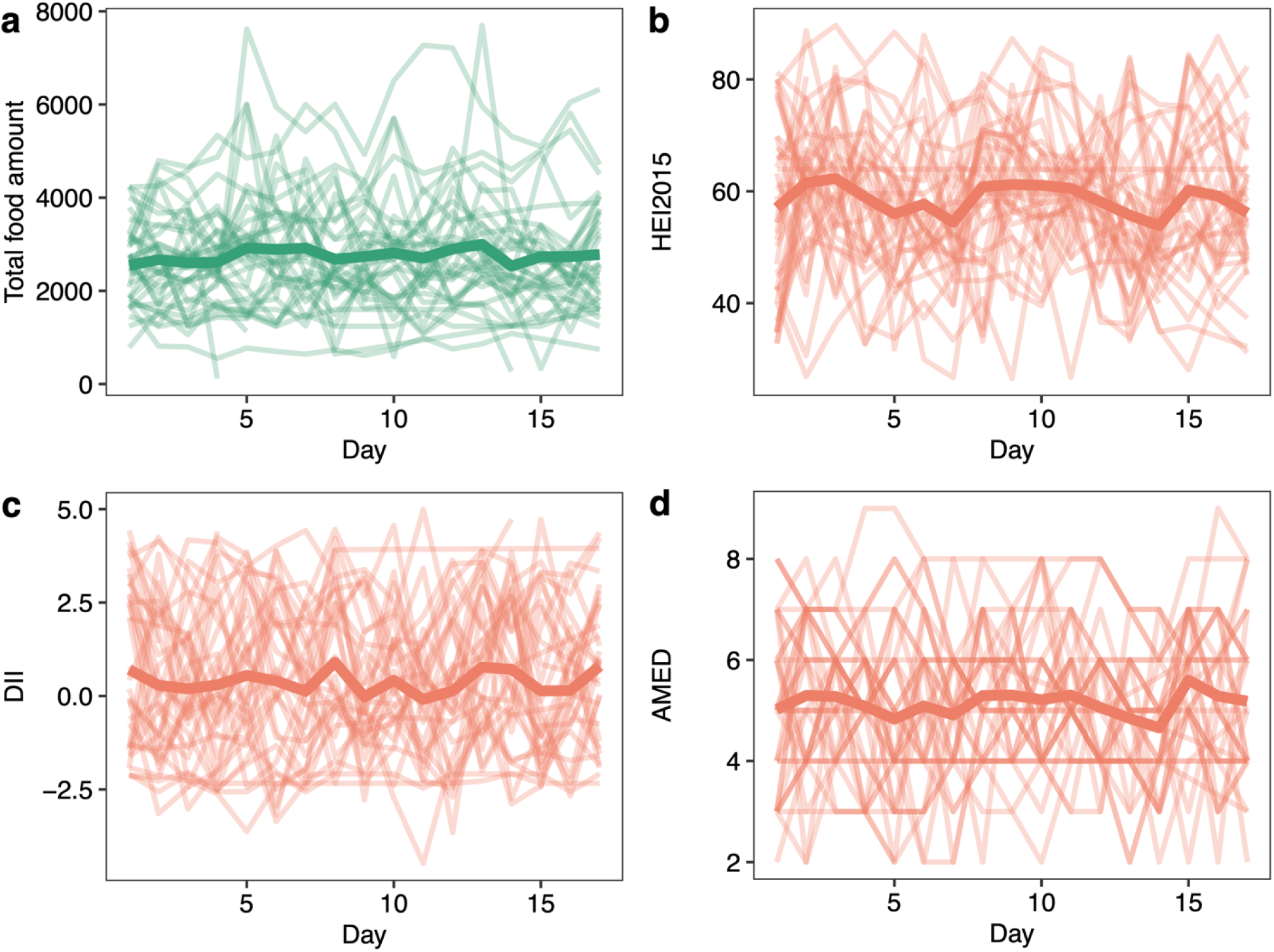
Consumption patterns and diet score in DMAS. **a**, Total food grams of each subject across 17 days. A thin line represents a subject, and the thick line represents the mean of all 30 subjects. **b-d**, HEI2015 (b), DII (c), and AMED (d) of each subject across 17 days. A thin line represents a subject, and the thick line represents the mean of all 30 subjects.

**Figure S2:**
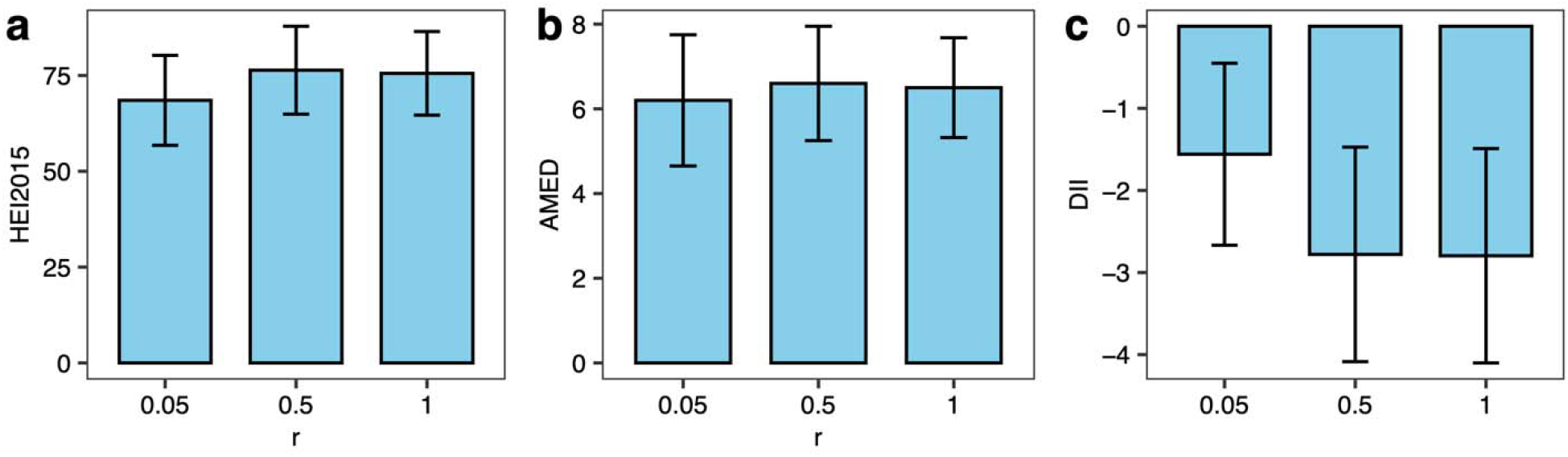
Relationship between diet score and parameter. Each bar represents the mean diet score, and the error bar indicates the standard deviation for HEI-2015 (a), AMED (b), and DII (c) among 10 randomly selected subjects in the DMAS cohort.

